# Therapeutic potential of macrophage colony-stimulating factor (CSF1) in chronic liver disease

**DOI:** 10.1101/2021.11.07.467663

**Authors:** Sahar Keshvari, Berit Genz, Ngari Teakle, Melanie Caruso, Michelle F. Cestari, Omkar L. Patkar, Brian WC Tse, Kamil A Sokolowski, Hilmar Ebersbach, Julia Jascur, Kelli P. A. MacDonald, Gregory Miller, Grant A. Ramm, Allison R. Pettit, Andrew D. Clouston, Elizabeth E. Powell, David A. Hume, Katharine M. Irvine

## Abstract

Resident and recruited macrophages control the development and proliferation of the liver. We showed previously in multiple species that treatment with a macrophage colony stimulating factor (CSF1)-Fc fusion protein initiated hepatocyte proliferation and promoted repair in models of acute hepatic injury in mice. Here we investigated the impact of CSF1-Fc on resolution of advanced fibrosis and liver regeneration, utilizing a non-resolving toxin-induced model of chronic liver injury and fibrosis in C57BL/6J mice. Co-administration of CSF1-Fc with exposure to thioacetamide (TAA) exacerbated inflammation consistent with monocyte contributions to initiation of pathology. After removal of TAA, either acute or chronic CSF1-Fc treatment promoted liver growth, prevented progression and promoted resolution of fibrosis. Acute CSF1-Fc treatment was also anti-fibrotic and pro-regenerative in a model of partial hepatectomy in mice with established fibrosis. The beneficial impacts of CSF1-Fc treatment were associated with monocyte-macrophage recruitment and increased expression of remodeling enzymes and growth factors. These studies indicate that CSF1-dependent macrophages contribute to both initiation and resolution of fibrotic injury and that CSF1-Fc has therapeutic potential in human liver disease.

**Summary statement:** Macrophages contribute to both progression and resolution of chronic tissue injury and fibrogenesis. Administration of a macrophage growth factor promoted liver regeneration and resolution of advanced liver fibrosis in mice.

## Introduction

Fibrosis is a physiological response to acute and chronic tissue injury. In the developed world 45% of all-cause mortality may be attributable to fibrotic disorders (Wynn, 2008). Chronic liver disease (CLD) and associated liver fibrosis, cirrhosis and its complications and hepatocellular carcinoma (HCC) affect more than 1.5 billion people and cause more than 2 million deaths each year (Asrani et al., 2019; Moon et al., 2020). Liver injury of any etiology causes an inflammatory response leading to myofibroblast activation and collagen deposition (Wynn, 2008). Where the injury persists, ongoing extracellular matrix (ECM) deposition leads to disruption of the liver architecture, eventually leading to portal hypertension and loss of liver function characteristic of cirrhosis. Patients with cirrhosis are at high risk of life-threatening complications and HCC. Despite an active clinical trial pipeline (Lemoinne and Friedman, 2019), no anti-fibrotic therapies are available.

Liver fibrogenesis is a dynamic process that can be modulated by preventing progression or promoting resolution. Fibrosis may reverse in patients receiving successful antiviral therapy and in abstinent alcohol-induced cirrhosis patients (D’Ambrosio et al., 2012; Marcellin et al., 2013; Schuppan et al., 2018) but many patients with advanced fibrosis due to viral hepatitis or alcohol-induced cirrhosis progress despite removal of the primary stimulus (Schuppan et al., 2018). Fibrosis also complicates surgery in CLD patients (for example to remove a tumour) since it can also impair regeneration (Hackl et al., 2016; Krenzien et al., 2018).

Macrophages and monocytes contribute to both disease progression and resolution in chronic liver disease (Duffield et al., 2005; Irvine et al., 2019; Kisseleva and Brenner, 2021; Tacke, 2017). Inhibition of some macrophage functions (e.g. Wnt secretion, autophagy, phagocytosis) during disease progression can exacerbate fibrosis (Irvine et al., 2015; Lodder et al., 2015; Perugorria et al., 2018; Wan et al., 2020). During fibrosis resolution macrophages may transition to a pro-repair phenotype, clearing damaged tissue, dampening inflammation and fibroblast activation, and producing growth and matrix remodelling factors that reduce fibrous tissue and restore liver architecture (Ramachandran et al., 2012). However, in contrast to the significant interest in macrophages as therapeutic targets to limit inflammation (Tacke, 2017), few therapeutic approaches to promote macrophage pro-regenerative functions have been little explored (Barcena et al., 2019; Han et al., 2019; Wan et al., 2020).

Signaling through the macrophage colony-stimulating factor receptor (CSF1R) drives monocyte differentiation, proliferation and function (Hume et al., 2019). To test potential therapeutic applications in tissue repair, we generated a porcine CSF1-Fc fusion protein that has an extended circulating half-life compared to the native protein. CSF1-Fc treatment promoted hepatocyte proliferation and liver growth in healthy mice, rats and pigs (Gow et al., 2014; Irvine et al., 2020; Sauter et al., 2016). Acute CSF1-Fc treatment increased liver macrophage content through both CCR2-dependent monocyte infiltration and resident macrophage proliferation (Gow et al., 2014; Stutchfield et al., 2015). CSF1R is expressed exclusively in cells of the macrophage lineage (Grabert et al., 2020) so the effect on hepatocytes must reflect indirect impacts of expansion of liver monocyte-macrophage populations. CSF1-Fc improved healing in paracetamol-induced acute liver failure in mice, increased regeneration of healthy liver following partial hepatectomy (Stutchfield et al., 2015) and promoted recovery from ischemia reperfusion injury in fibrotic liver (Konishi et al., 2020). These data suggest CSF1 has therapeutic potential in liver disease. Infusion of CSF1-differentiated bone-marrow derived macrophages showed promise in murine models and is currently being tested in the clinic (Dwyer et al., 2021; Thomas et al., 2011). However, some of the earliest studies of CSF1 in disease models also showed the potential for exacerbation of inflammatory pathology (Hume and MacDonald, 2012), and macrophage depletion with anti-CSF1R or anti-CSF1 was previously shown to ameliorate development of toxin-induced liver fibrosis in mice (Mehal et al., 2001; Seifert et al., 2015). Here we tested the therapeutic potential of CSF1-Fc and the role of macrophages in resolution of liver fibrosis in a non-resolving model of chronic inflammatory liver injury.

## Results

### Chronic CSF1-Fc treatment initiates resolution of established fibrosis

The model we have chosen is chronic exposure to thioacetamide (TAA) in the drinking water. Both TAA and the more-widely employed carbon tetrachloride (CCl4) model a sustained sterile injury to hepatocytes (similar to alcohol, for example), leading to progressive inflammation and fibrogenesis. The CCL4 model has previously been used to investigate macrophage contributions to fibrosis progression and regression (Duffield et al., 2005; Ramachandran et al., 2012; Thomas et al., 2011) but unlike TAA-induced injury, CCl4 hepatotoxicity and fibrosis rapidly and spontaneously resolves (Liu et al., 2020; Ramachandran et al., 2012).

Previous studies showed that acute porcine (P)-CSF1-Fc treatment (4 daily injections of 1 mg/kg) led to a monocytosis, monocyte-macrophage accumulation in liver, hepatocyte proliferation and liver growth (Gow et al., 2014; Irvine et al., 2020; Sauter et al., 2016). In a pilot experiment, mice treated twice weekly with 1 mg/kg P-CSF1-Fc commencing at the same time as TAA administration experienced rapid onset of toxicity and did not survive past day 10-14. The CSF1-Fc treatment greatly exacerbated the hepatic pericentral inflammatory infiltration that commences within 1 week of TAA exposure (Melino et al., 2016). This was associated with alpha smooth muscle actin (αSMA) expression but not collagen deposition, nor extensive hepatocyte necrosis (**Fig. S1**). These observations indicate that TAA does not prevent the response to CSF1-Fc, and not surprisingly, increasing monocyte recruitment in a setting of ongoing acute injury associated with generation of damage-associated molecular patterns (DAMPs) promotes pathology.

To model treatment of established liver disease in humans following removal of the primary stimulus, we tested the impact of CSF1-Fc treatment on liver fibrosis resolution after TAA cessation. In addition to P-CSF1-Fc, these experiments utilized a human CSF1-mouse Fc conjugate (HM-CSF1-Fc), which was developed by Novartis (Switzerland) for evaluation in preclinical models. HM-CSF1-Fc was utilised at 5 mg/kg as initial studies demonstrated this dose induced a comparable biological response to 1 mg/kg P-CSF1-Fc (**Fig. S2**, see Materials and Methods for further details of CSF1-Fc reagents). Female mice were administered TAA in drinking water for 8 weeks or provided normal drinking water, followed by TAA withdrawal and twice-weekly treatment with CSF1-Fc or saline for 4 weeks, with sacrifice 1 day following the final dose. Additional groups of mice were left to recover for 4 weeks (**Fig. 1A**). Blood monocyte count was no longer elevated in CSF1-Fc-treated mice by this timepoint (**Fig. 1B**), but liver monocyte and macrophage content was increased reflected by increased mRNA expression of the monocyte chemokine *Ccl2* and receptor *Ccr2*, as well as *Adgre1* (encoding the monocyte/macrophage marker F4/80), particularly in TAA-treated mice (**Fig. 1C-E**). Flow cytometry analysis of disaggregated liver confirmed the increases in both monocyte-derived (F4/80^Hi^/TIM4^−^) and resident (F4/80^Hi^/TIM4^+^) liver macrophages and the infiltration of monocytes (F4/80^Low^/Cd11b^Hi^) in CSF1-Fc-treated mice (**Fig. S3A-C**). CSF1 drives the maturation of Ly6C^Hi^ into Ly6C^Low^ monocytes, which have previously been associated with resolution of liver fibrosis (Ramachandran et al., 2012), and their subsequent differentiation into macrophages. Consistent with this, a higher proportion of liver macrophages in CSF1-Fc-treated livers were monocyte-derived (TIM4^−^) compared to saline-treated mice (**Fig. S3B**). Together these data demonstrate that CSF1-Fc promotes recruitment and maturation of monocytes in the liver regardless of prior injury in response to TAA exposure. The increase in hepatic monocyte/macrophages was reversible and had returned to baseline 4 weeks after the cessation of CSF1-Fc treatment (**Fig. 1C-E**).

**Fig. 1.**
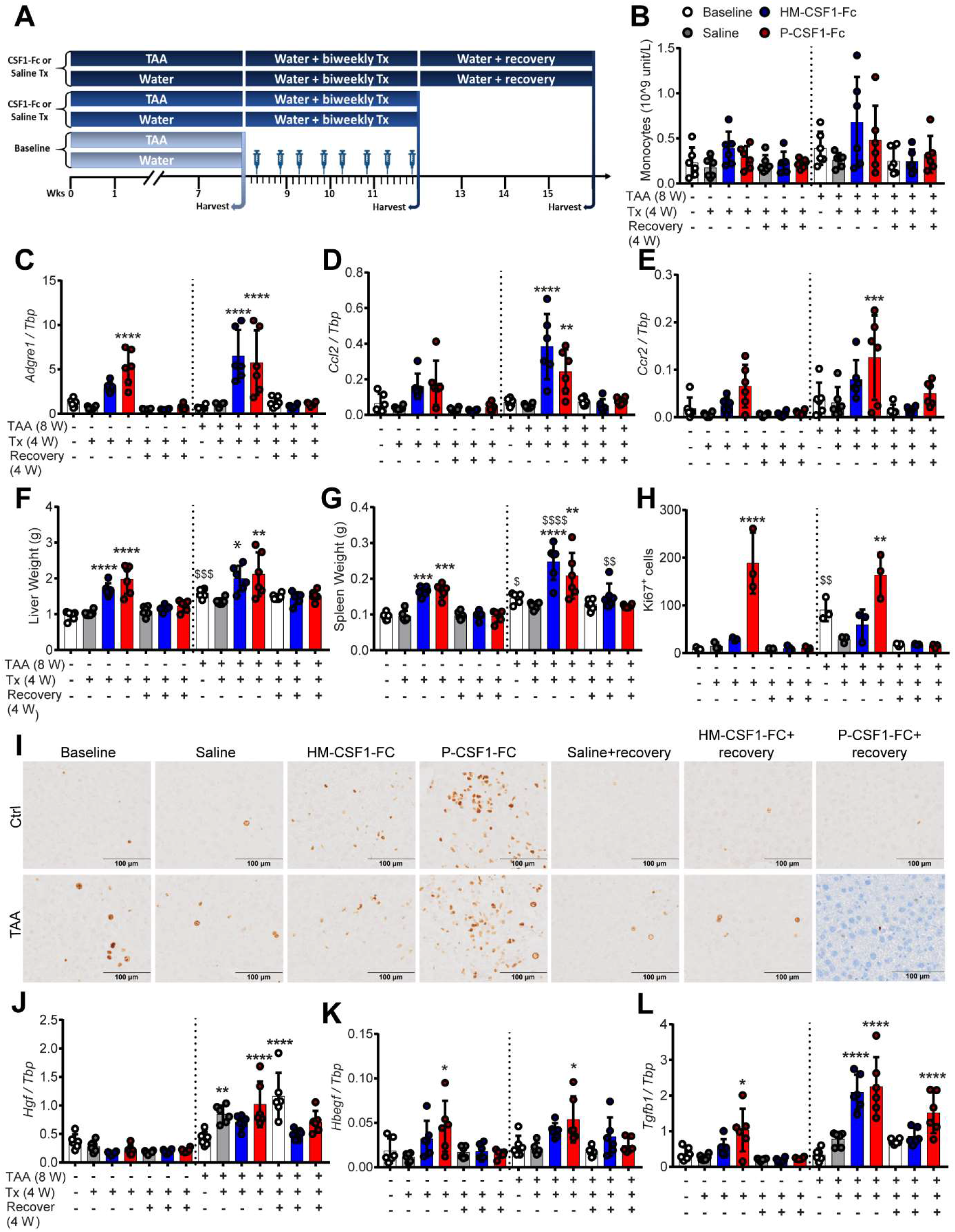
Chronic CSF1-Fc-induced expansion of liver macrophages drives growth of chronically injured liver. (A) Male mice were administered TAA for 8 weeks, prior to 4 weeks treatment with P-CSF-Fc, HM-CSF1-Fc or saline. (B) Blood monocyte count. Whole liver expression of (C) *Adgre1*, (D) *Ccl2*, and (E) Ccr2. (F) Liver and (G) spleen weight. (H,I) Liver Ki67+ cells. Whole liver expression of (J) *Hgf*, (K) *Hbegf*, and (L) *Tgfb1*. One-Way ANOVA with multiple comparisons: *p<0.05, **p<0.01, ***p<0.001, ****p<0.0001 comparing to baseline in the same group. $ p<0.05, $$ p<0.01, $$$ p<0.001, $$$$ p<0.0001 comparing the same treatment between groups (n=6 per group with 3 animals per group randomly selected for IHC image analysis).

CSF1-Fc treatment increased the size of both the liver and spleen in control mice as reported previously. This response was also observed in TAA-exposed mice and returned to baseline 4 weeks after cessation of treatment (**Fig. 1F,G**). CSF1-Fc-induced liver growth was associated with proliferation of non-parenchymal cells (**Fig. 1H,I, Fig. S4A**). Among liver mitogens, previous analysis revealed that acute CSF1-Fc treatment induced expression of heparin binding EGF-like growth factor (*Hbegf*) in healthy mice, but did not affect *Egf*, hepatocyte growth factor (*Hgf*), transforming growth factor alpha or amphiregulin (Gow et al., 2014). *Hgf* mRNA increased upon TAA cessation, but was not further induced by CSF1-Fc (**Fig. 1J**). Both CSF1-Fc reagents induced *Hbegf* regardless of liver injury, as well as transforming growth factor (*Tgfb1*), an established mediator of fibroblast activation and fibrosis that also drives the resident liver macrophage transcriptional programme (Sakai et al., 2019) (**Fig. 1K,L**).

Eight weeks TAA administration induced bridging fibrosis/cirrhosis (**Fig. 2A**). Histological inflammation (not shown) and expression of the major fibrillar collagens *Col1A1* and *Col3A1* (**Fig. 2B, Fig. S4B**) was reduced 4 weeks after TAA cessation. Expression of *Acta2* (encoding smooth muscle alpha-actin; αSMA), generally considered a marker of myofibroblast/stellate cell activation, was variable and was not significantly elevated by TAA or CSF1-Fc in this chronic model (**Fig. 2C**). Acute CSF1-Fc treatment was previously reported to induce hepatic expression of inflammatory cytokines including *Il6* and *Tnf* (Stutchfield et al., 2021). *Il6* was significantly upregulated by chronic CSF1-Fc treatment in healthy but not TAA-treated liver, whilst *Tnf* was upregulated during TAA regression and not further regulated by CSF1-Fc (**Fig. S4C,D**).

**Fig. 2.**
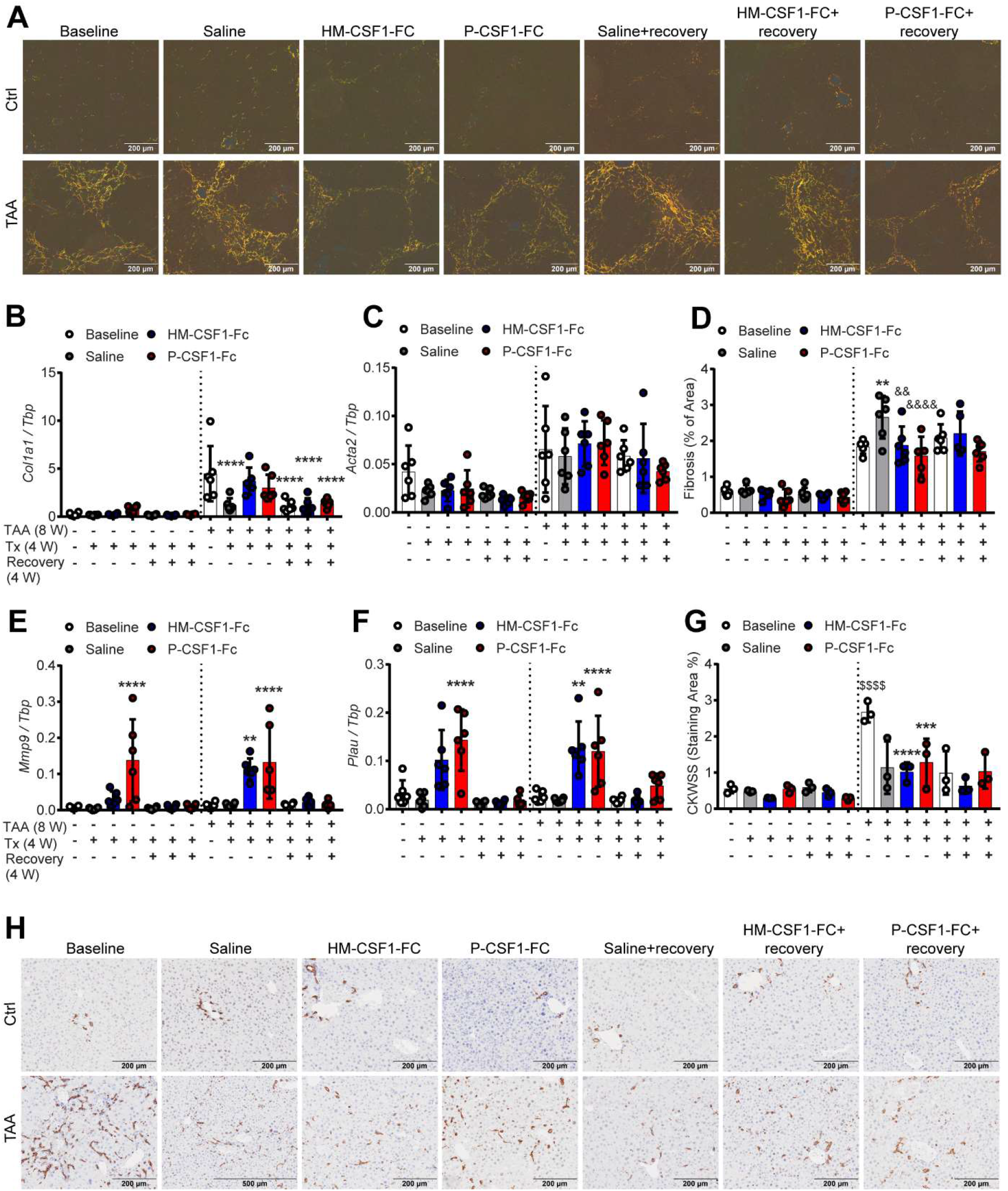
Chronic CSF1-Fc treatment initiates resolution of established fibrosis. Experimental design as in Fig. 1A. (A) Collagen was assessed by picrosirius red staining visualised under polarised light and (D) quantified by image analysis. Expression of (B) *Colla1*, (C) *Acta2*, (E) *Mmp9 and* (F) *Plau* in whole liver was quantified by RT-PCR. (H) Hepatic progenitor cell activation was assessed by CKWSS immunohistochemistry and (G) quantified by image analysis. One-Way ANOVA with multiple comparison, **p<0.01, ***p<0.001, ****p<0.0001 comparing to baseline in the same group. && p<0.01, &&&& p<0.0001 comparing to Saline in same group.$$$$ p<0.0001 comparing the same treatment between groups (n=6 per group with 3 animals per group randomly selected for IHC image analysis).

Consistent with previous reports in the TAA model (Delire et al., 2015; Popov et al., 2011), fibrosis did not spontaneously regress following removal of the stimulus. In fact there was a further small increase in fibrotic area 4 weeks after TAA cessation (**Fig. 2D)**, which was prevented by CSF1-Fc treatment. CSF1-Fc treatment induced expression of matrix remodeling genes including *Mmp9*, *Mmp13* and *Plau* (encoding urokinase plasminogen activator, uPA) (**Fig. 2E,F, Fig. S4E**). *Mmp9* is highly expressed by resident and recruited monocyte-derived macrophages in multiple mouse organs including liver (Summers et al., 2020). *Plau* is a CSF1R target gene in macrophages (Stacey et al., 1995). As in all chronic liver injury models, TAA induced prominent hepatic progenitor cell (HPC) activation, identified by wide-spectrum keratin (CKWSS) staining (Melino et al., 2016). Unlike the fibrosis, HPC activation largely resolved spontaneously by 4 weeks after TAA cessation and CSF1-Fc treatment had minimal impact (**Fig. 2G,H**).

To determine whether a less frequent CSF1-Fc treatment regime would effectively remodel the fibrotic scar we investigated the impact of once weekly treatment on fibrosis regression. This regimen was effective in a fracture healing model (Batoon et al., 2021). Female mice were administered TAA for 8 weeks, followed by treatment with CSF1-Fc for up to 8 weeks (mice were sacrificed one week following the final dose). Weekly CSF1-Fc treatment did not significantly ameliorate fibrosis or persistently increase liver weight and the impact on macrophage and matrix remodeling gene expression at the 4 week timepoint was not sustained (**Fig. S5A-E**). This led us to question whether CSF1-Fc from other species induces an anti-drug antibody response that may limit the impact of treatment over time. Indeed, there was a significant anti-CSF1-Fc response in mice treated with either CSF1-Fc reagent (**Fig. S5F,G)**. In overview, these studies indicate CSF1-Fc treatment has potential to initiate liver fibrosis resolution, but evaluation of chronic treatment regimens with existing reagents in mice is compromised by an anti-drug response.

### Acute CSF1-Fc treatment is sufficient to eliminate established fibrosis

Previous studies of CSF1-Fc treatment in acute liver injury models utilised 4 successive daily injections (Stutchfield et al., 2015). To test an acute regime, male and female mice were treated with TAA for 8 weeks, followed by 4 daily injections of CSF1-Fc or saline prior to sacrifice on day 5, or recovery to day 14 (**Fig. 3A**). This acute regime induced monocytosis and increased liver and spleen weight as expected (**Fig. 3B-D**). Here we showed that this impact is rapidly reversible and had resolved by day 14. Acute CSF1-Fc induced *Adgre1, Ccl2* and *Ccr2* mRNA in the liver, consistent with an increase in monocytes and macrophages (**Fig. 3E-G**). As in the chronic treatment, CSF1-Fc transiently induced *Tgfb1*, but not *Il6* and *Tnf* (**Fig. 3H-J**). CSF1-Fc-induced liver growth was associated with an increase in Ki67+ parenchymal and non-parenchymal cells, which normalized by day 14 (**Fig. 3K-M**). No differences in *Hgf* or *Hbegf* expression were observed (**Fig. 3N,O**). CSF1-Fc did not increase CKWSS staining area, suggesting HPC do not contribute to the large increase in proliferating non-parenchymal cells or to liver growth (**Fig. 4A-B**).

**Fig. 3.**
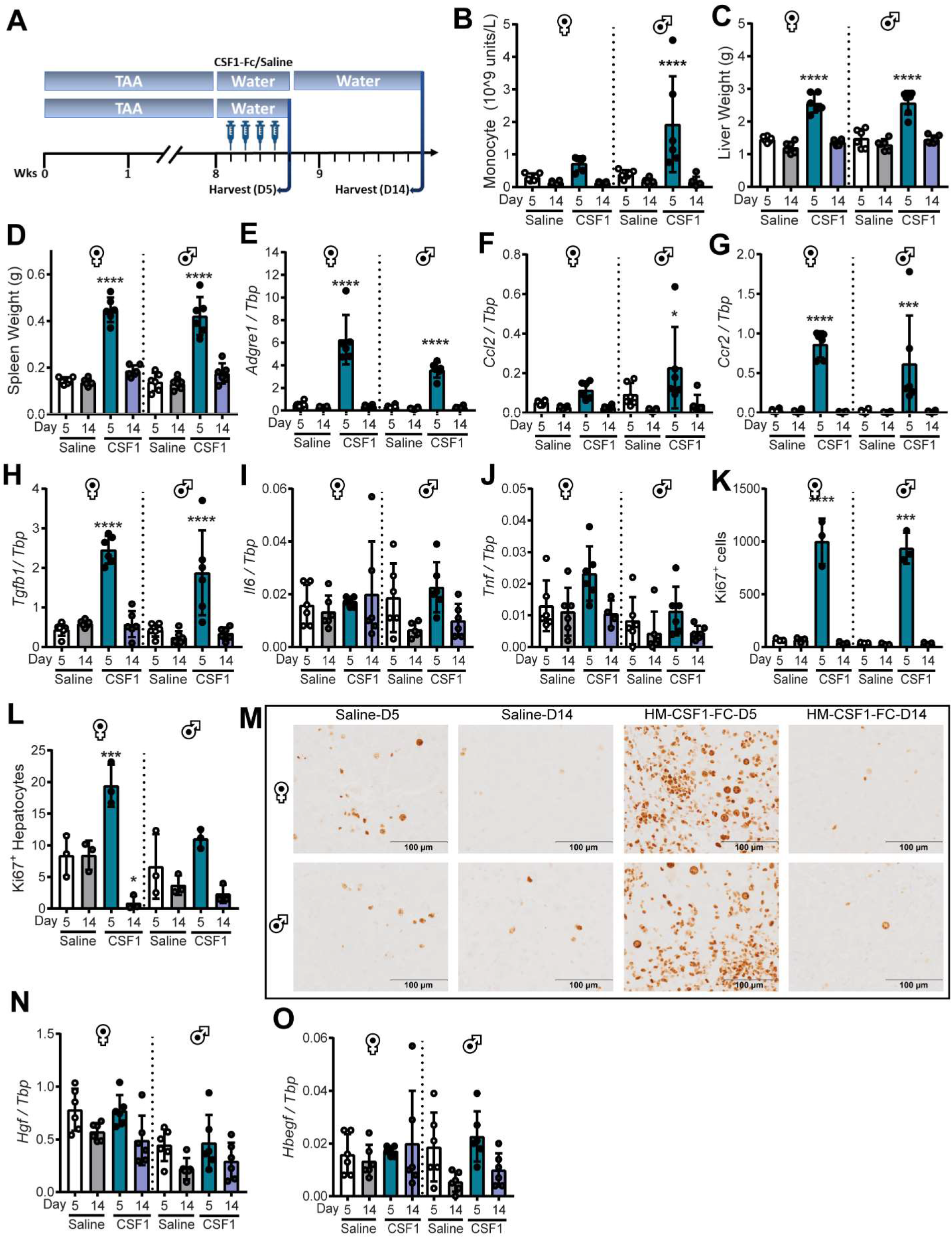
Acute CSF1-Fc-induced expansion of liver macrophages drives growth of chronically injured liver. (A) Male and female mice were administered TAA for 8 weeks, prior to daily treatment with HM-CSF1-Fc or saline for 4 days, and sacrificed on day 5 or 14. (B) Blood monocyte count. (C) Liver and (D) spleen weight. Whole liver expression of (E) *Adgre1*, (F) *Ccl2*, (G) Ccr2, (H) *Tgfb1*, (I) *Il6*, (J) *Tnf*, (N) *Hgf* and (O) *Hbegf*. (K, L) Liver Ki67+ cells and (L) hepatocytes. One-Way ANOVA with multiple comparison, *p<0.05, **p<0.01, ***p<0.001, ****p<0.0001 comparing to saline day 5 in the same group (n=6 per group with 3 animals per group randomly selected for IHC image analysis).

**Fig. 4.**
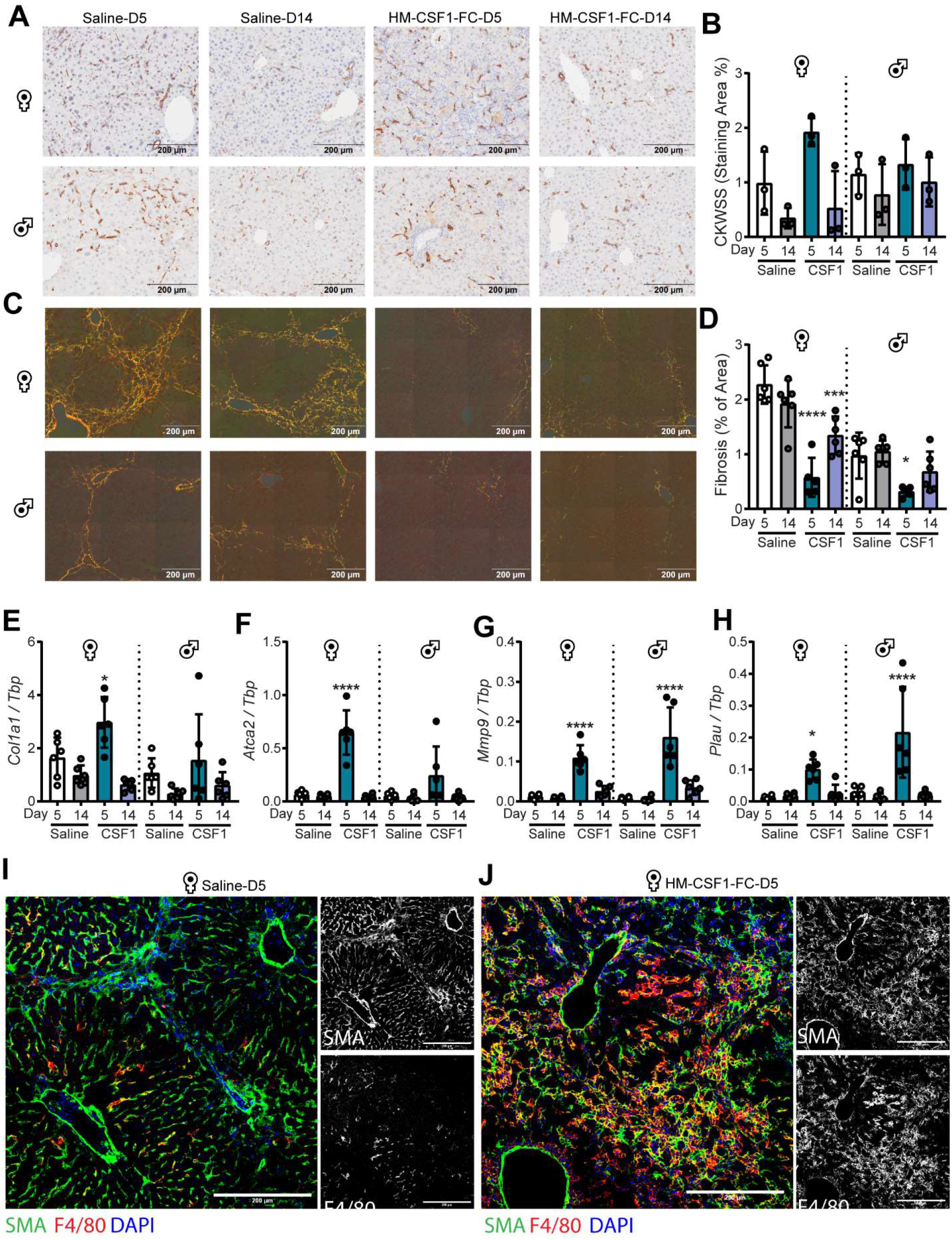
Acute CSF1-Fc treatment resolves established fibrosis. Experimental design as in Fig. 3A. Hepatic (A,B) progenitor cell activation was assessed by CKWSS immunohistochemistry. (C,D) Collagen was assessed by picrosirius red staining visualised under polarised light. Whole liver expression of (E) *Col1a1*, (F) *Acta2*, (G) *Mmp9* and (H) *Plau*. Representative SMA and F4/80 staining in (I) saline and (J) CSF1-Fc treated liver (1 representative animal per group). One-Way ANOVA with multiple comparison, *p<0.05, **p<0.01, ***p<0.001, ****p<0.0001 comparing to saline day 5 in the same group (n=6 per group with 3 animals per group randomly selected for IHC image analysis).

Acute CSF1-Fc treatment promoted a significant reduction in fibrotic area, especially in the females which were more severely affected (**Fig. 4C,D**). This effect was partly reversed by day 14 post-treatment. Fibrosis regression was associated with transient increases in *Mmp9* and *Plau* expression on day 5, which also returned to baseline by day 14 (**Fig. 4G,H)**. Surprisingly, CSF1-Fc treatment also produced a transient increase in *Acta2* mRNA encoding αSMA (**Fig. 4E**), whereas *Colla1* was unaffected (**Fig. 4F**). In saline-treated mice αSMA was restricted to sinusoidal myofibroblasts. By contrast, in CSF1-Fc treated mice increased αSMA expression co-localised with F4/80^+^ macrophages (**Fig. 4I,J**).

### CSF1-Fc treatment reduces fibrosis and improves fibrotic liver regeneration after partial hepatectomy

Liver regenerative capacity in patients with advanced fibrosis is compromised, which limits surgical intervention (Hackl et al., 2016; Krenzien et al., 2018). To model application of acute CSF1-Fc treatment in this indication, we established fibrosis using 8 weeks TAA treatment, then performed a 50% partial hepatectomy (PHx). Mice were treated with CSF1-Fc or saline for 2 days prior and 2 days post-surgery, followed by sacrifice on day 3 (**Fig. 5A**). Mice with both healthy and fibrotic liver treated with CSF1-Fc gained weight post-surgery more rapidly and had increased liver and spleen mass on day 3 (**Fig. 5B-D**). CSF1-Fc treatment increased circulating monocytes, as well as liver *Adgre1*, *Ccl2* and *Ccr2* expression (**Fig. 5E-H**). PHx-induced liver regrowth was associated with an increase in proliferative (Ki67+) hepatocytes and non-parenchymal cells in the remnant liver, which was partly compromised in fibrotic livers (**Fig. 5J-L**). CSF1-Fc treatment increased Ki67+ non-parenchymal cells but did not further increase Ki67+ hepatocytes at this time point, and did not overcome the deficit in the fibrotic livers (**Fig. 5K,L**). Nevertheless, CSF1-Fc treated mice had substantial increases in mRNA encoding *Hbegf*, *Tgfb1*, *Il6* and *Tnf* (**Fig. 5O-R**) each of which could contribute to hepatic growth (Kiso et al., 2003; Michalopoulos and Bhushan, 2021). *Hgf* and *Brg1*, a chromatin remodelling gene involved in liver regeneration (Wang et al., 2019), were more highly expressed following PHx in fibrotic liver compared to healthy liver (**Fig. 5I,S**). Neither PHx nor CSF1-Fc affected HPC abundance (**Fig. 5J,N**). CSF1-Fc treatment almost completely eliminated fibrosis in this model (**Fig. 5J,M**). This was associated with reduced hepatic *Col3a1* but not *Colla1* expression (**Fig. 6A,B**). The CSF1-Fc-induced fibrosis resolution was again associated with increases in *Mmp9* and *Plau* (**Fig. 6C,D**).

**Fig. 5.**
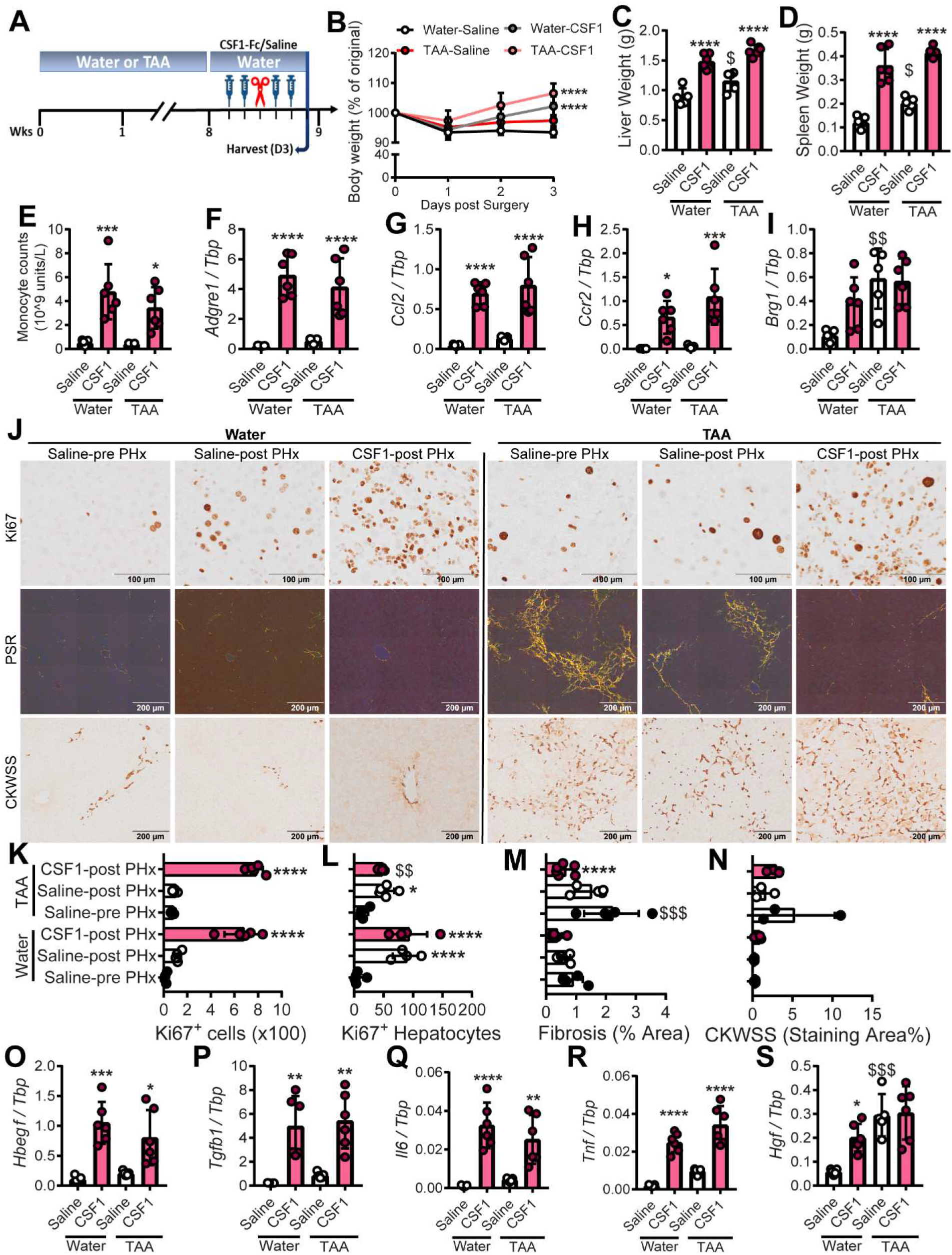
CSF1-Fc promotes liver regrowth and fibrosis resolution post-resection. (A) Male mice were administered TAA/normal water for 8 weeks, treated with P-CSF1-Fc or saline pre- and post-50% hepatectomy; sacrifice on day 3. (B) Body, (C) liver and (D) spleen weight. (E) Blood monocytes. Whole liver expression of (F) *Adgre1*, (G) *Ccl2*, (H) Ccr2, (I)*Brg1*, (O) *Hbegf*, (P) *Tgfb1*, (Q) *I16*, (R) *Tnf* and (S) *Hgf*. Ki67+ cells (J,K,L). Collagen (picrosirius red, J,M. HPC (CKWSS, J,N). One-Way ANOVA with multiple comparison, *p<0.05, **p<0.01, ***p<0.001, ****p<0.0001 comparing to saline in the same group. $p<0.05, $$p<0.01, $$$p<0.001, $$$$p<0.0001 comparing the same treatment between groups (n=3 per group for water treatment, and 6 per group for TAA treatment with 3 animals per group randomly selected for IHC image analysis).

**Fig. 6.**
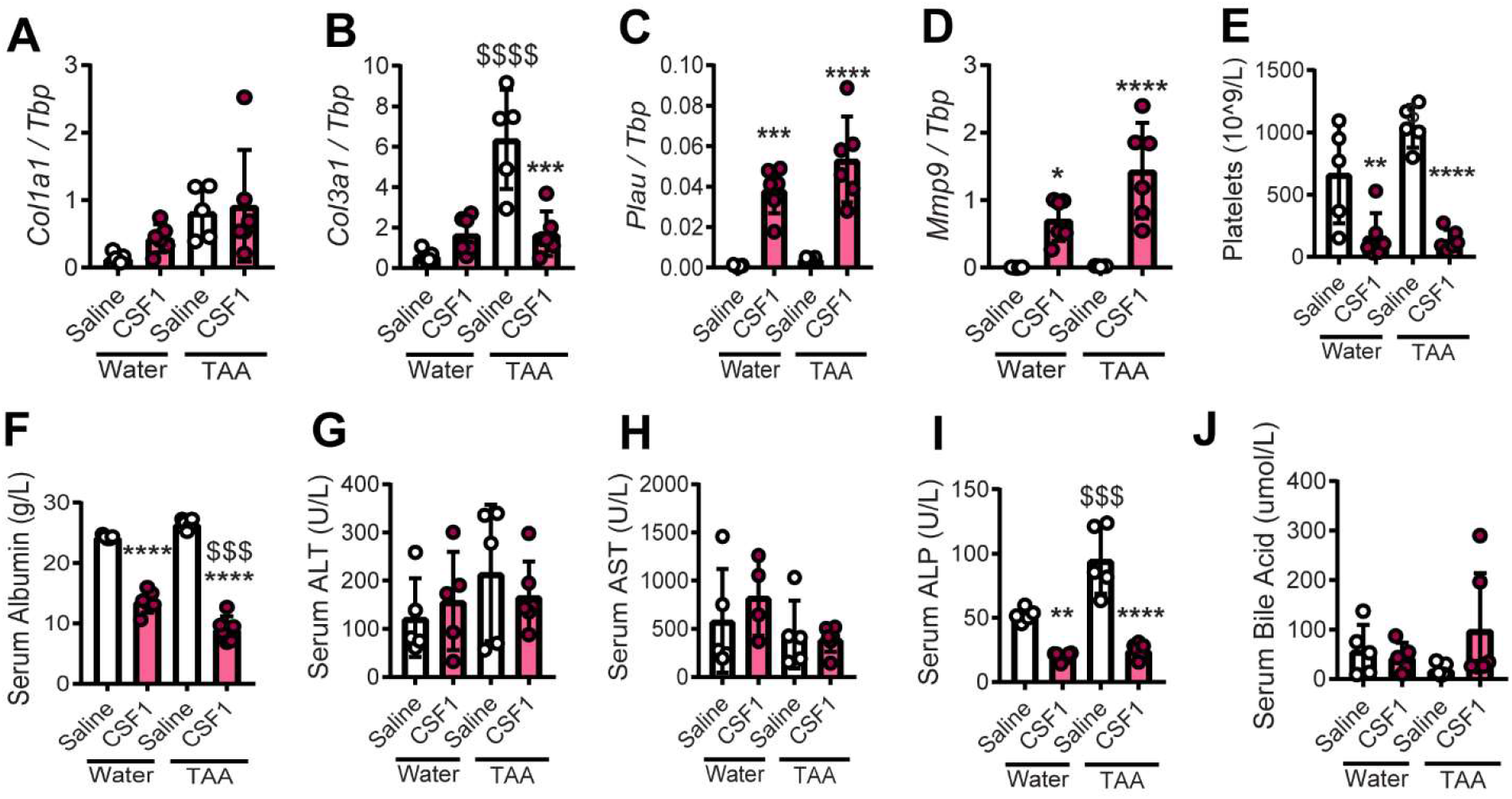
Impacts of acute CSF1-Fc treatment on hepatic gene expression and serum biochemistry post-resection. Experimental design as in Fig. 5A. Whole liver expression of (A) *Col1a1*, (B) *Col3a1*, (C) *Plau*, and (D) *Mmp9*. (E) Circulating platelets, (F) Serum albumin, (G) ALT, (H) AST, (I) ALP and (J) bile acids. One-Way ANOVA with multiple comparison, *p<0.05, **p<0.01, ***p<0.001, ****p<0.0001 comparing to saline in the same group. $ p<0.05, $$ p<0.01, $$$ p<0.001, $$$$ p<0.0001 comparing the same treatment between groups (n=3 per group for water treatment and 6 per group for TAA treatment).

One consequence of CSF1 treatment in animals and humans (Baker and Levin, 1998),(Garnick and O’Reilly, 1989) is a reversible reduction in blood platelets. We observed an extended clotting time, lower platelet count (**Fig. 6E**), reduced serum albumin (**Fig. 6F**), and apparent ascites in CSF1-Fc-treated mice that had undergone PHx. Serum liver enzymes (ALT and AST) did not indicate increased hepatocellular injury; indeed ALP was significantly reduced by CSF1-Fc treatment, and there was no change in bile acids (**Fig. 6G-J**). Nevertheless, thrombocytopenia is of concern in CLD patients especially those undergoing surgery.

To determine whether these impacts of CSF1-Fc could be avoided whilst retaining desirable impacts on liver regeneration and fibrosis, we tested the efficacy of a sub-maximal dose of CSF1-Fc (“low-dose”). Male mice were administered TAA or normal drinking water for 12 weeks, treated with 1 mg/kg HM-CSF1-Fc or saline control for 2 days prior and 2 days post 50% Phx, and sacrificed 3 or 7 days post-surgery (**Fig. 7A**). Half of the mice in the 7-day recovery group were randomly assigned to undergo blood flow imaging on day 3 post-surgery. Although low-dose CSF1-Fc did not induce monocytosis in healthy mice (**Fig. S2B**) it did increase blood monocytes in TAA-exposed animals on day 3 post-surgery, which largely normalised by day 7 (**Fig. 7B**). Low dose CSF1-Fc treatment still accelerated body and liver weight gain post-surgery, increased spleen weight (**Fig. 7C-E**) and transiently induced hepatic *Adgre1*, *Ccl2* and *Ccr2* expression (**Fig. 7F-H**). Low dose CSF1-Fc induced *Hbegf*, and *Tgfb1*, but not *Hgf*, on day 3 post surgery (**Fig. 7I-K**) and greatly increased the number of Ki67+ proliferating non-parenchymal cells on day 3, (**Fig. 7L-O**). Low-dose CSF1-Fc still promoted resolution of hepatic fibrosis by day 3, which persisted to day 7 (**Fig. 7P,Q**). The treatment did not impact *Colla1* expression but induced sustained *Mmp9* elevation in both healthy and fibrotic liver, and transient induction of *Plau* (**Fig. 8A-C**). The low dose did not completely prevent the thrombocytopenia or reduced circulating albumin, but no ascites or clotting impairment were evident and these parameters were fully resolved by day 7 (**Fig. 8D,E**). Serum ALT and ALP were reduced in CSF1-Fc-treated mice (**Fig. 8F,G**). Doppler imaging showed decreased liver vascularisation in TAA-treated mice compared to healthy mice on day 3 post-PHx, which was reversed by low-dose CSF1-Fc treatment (**Fig. 8H,I**).

**Fig. 7.**
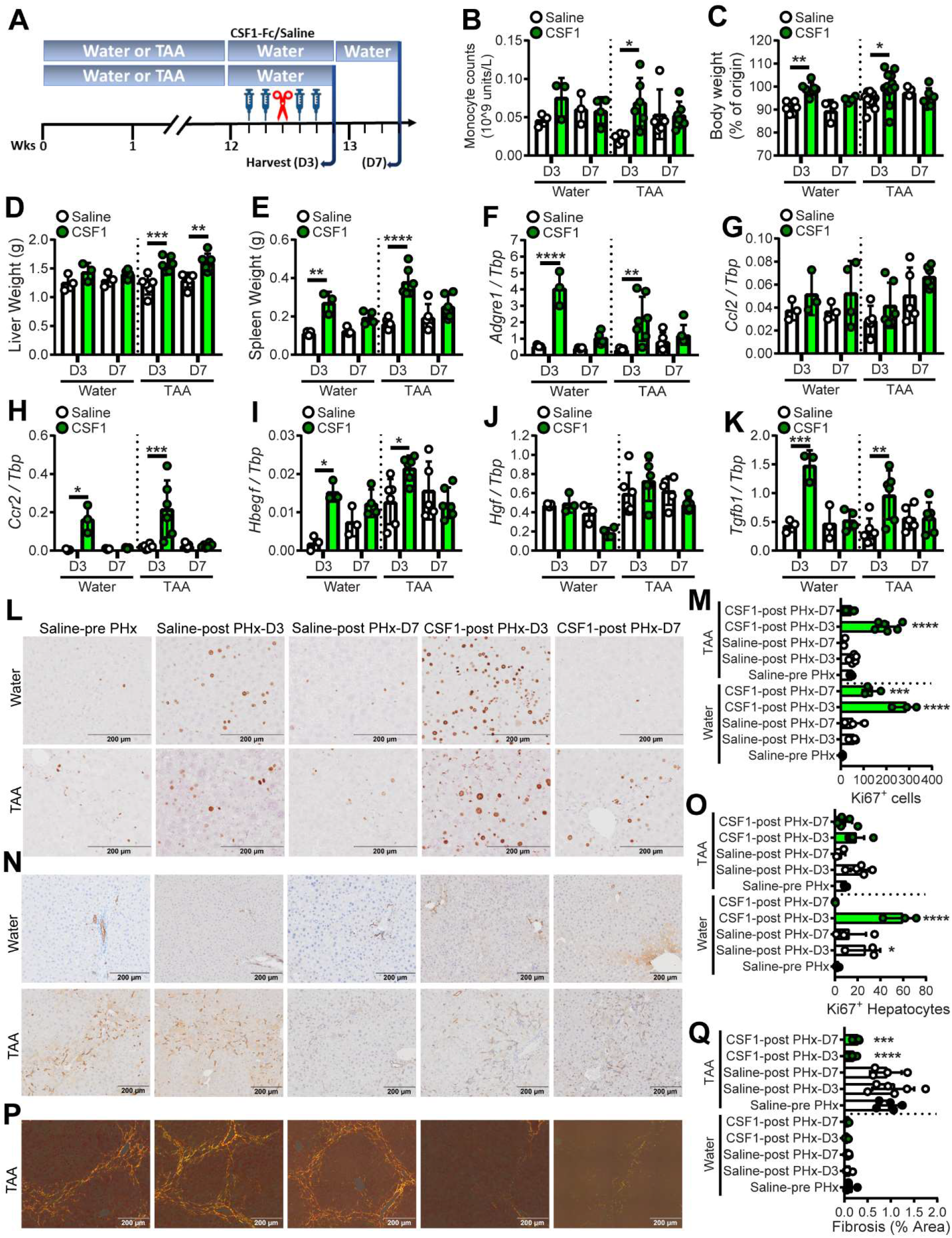
Low-dose CSF1-Fc treatment promotes liver regrowth and fibrosis resolution post-resection. (A) Male mice were administered TAA or normal water for 12 weeks, then treated with HM-CSF1-Fc or saline pre- and post-50% hepatectomy and sacrificed on day 3 or 7. (B) Blood monocyte count. (C) Body, (D) liver and (E) spleen weight. Whole liver expression of (F) *Adgre1*, (G) *Cc12*, (H) Ccr2, (I) *Hbegf*, (J) *Hgf*, and (K) *Tgfb1*. (L,M)Liver Ki67+ cells. (N,O) Liver HPC (CKWSS). (P,Q) Liver collagen (picrosirius red). One-Way ANOVA with multiple comparison, *p<0.05, **p<0.01, ***p<0.001, ****p<0.0001 comparing to saline pre PHx in the same group (n=3 per group for water treatment, and 6 per group for TAA treatment with 3 animals per group randomly selected for IHC image analysis).

**Figure 8.**
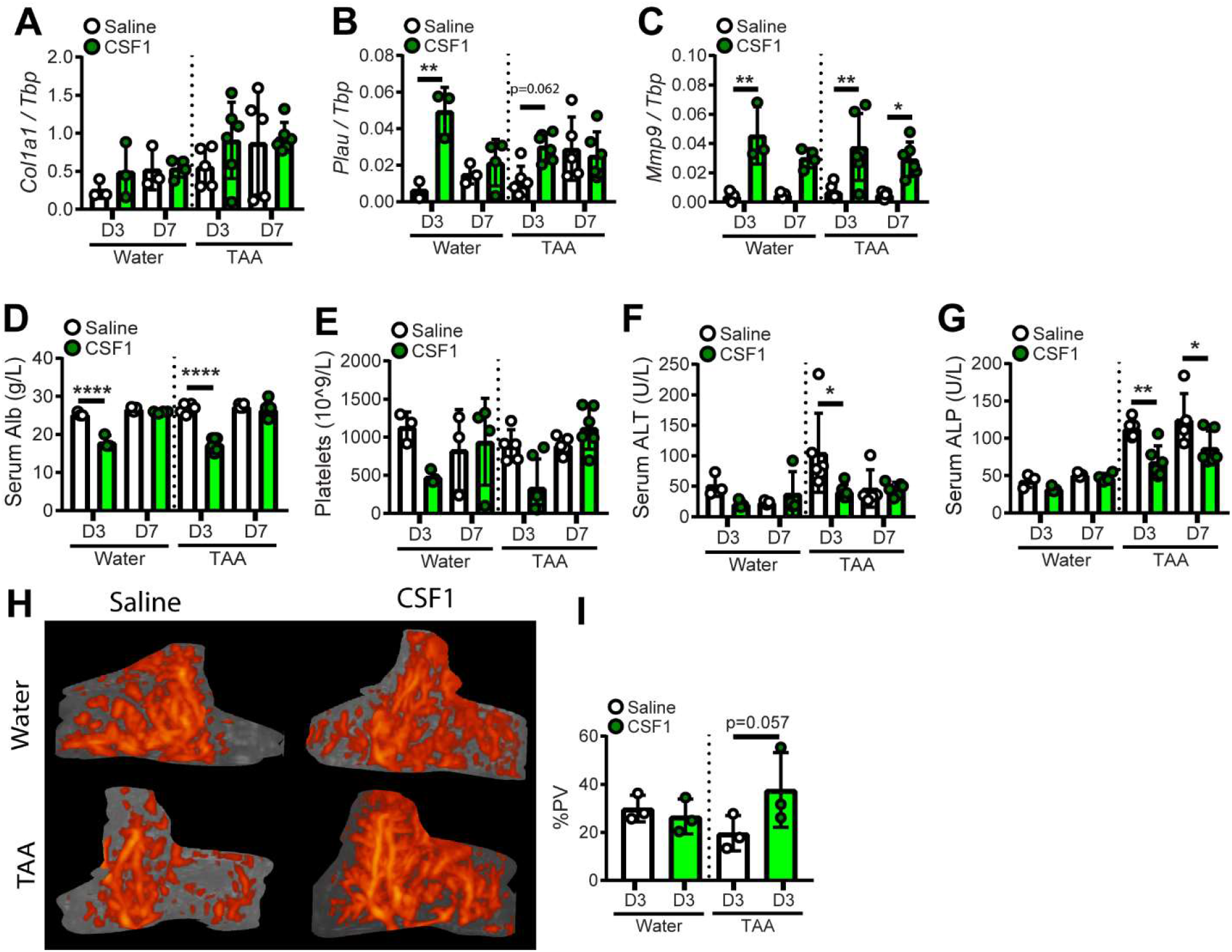
Impacts of low-dose CSF1-Fc treatment on hepatic gene expression, serum biochemistry and hepatic blood flow post-resection. Experimental design as in Fig. 7A. Whole liver expression of (A) *Colla1*, (B) *Plau*, and (C) *Mmp9*. Serum (D) albumin, circulating platelets (E), serum ALT (F) and ALP (G). (H,I) Hepatic blood flow was assessed by Power Doppler imaging. One-Way ANOVA with multiple comparison, *p<0.05, **p<0.01, ***p<0.001, ****p<0.0001 (n=3 per group for water treatment, and 6 per group for TAA treatment. n=3 randomly selected animals for Doppler imaging).

## Discussion

Resolution of fibrosis and liver regeneration post-surgical resection are unmet clinical needs. CSF1 and CSF1R are potential targets of anti-inflammatory treatments being developed by several pharmaceutical companies (Denny and Flanagan, 2021; Hamilton et al., 2016). CSF1 is a homeostatic growth factor that supports the trophic and matrix remodeling functions of macrophages during development and adult tissue maintenance, and which has been shown to promote liver growth in healthy mammals (Gow et al., 2014; Irvine et al., 2020; Sauter et al., 2016). Here we demonstrated that CSF1-Fc can promote hepatocyte proliferation and liver growth in chronically diseased liver, including after surgery. We also demonstrated that acute and chronic CSF1-Fc treatment significantly reduced the extensive fibrosis caused by long-term exposure to TAA, which does not resolve spontaneously. There is potential to investigate and optimize the chronic treatment regimes utilized here (Figure 1–2, Supplementary Figure 5), in which efficacy was likely compromised by the anti-drug antibody response, to achieve lasting resolution of fibrosis. We are currently developing a fully-orthologous mouse CSF1-Fc protein.

The therapeutic response to CSF1-Fc was associated with increased liver macrophages, a relative preponderance of the Ly6C^Low^ monocyte phenotype; and increased expression of matrix remodeling factors *Mmp9* and *Plau*. MMP9 overexpression promoted a pro-resolving macrophage phenotype and improved liver regeneration in cirrhotic mice (Melgar-Lesmes et al., 2018). Urokinase plasminogen activator (*Plau*) therapy ameliorated fibrosis in the CCL4 model via activation of latent metalloproteinases and HGF (Bueno et al., 2006; Meza-Rios et al., 2016; Salgado et al., 2000). Fibrosis regression occurred despite high expression of the canonical profibrogenic cytokine *Tgfb1* and, in some cases, the proinflammatory cytokines *Il6* and Tnf, which have frequently been shown to diminish in the resolution phase of disease. Given its pivotal role in the transcriptional programme driving liver macrophage differentiation (Sakai et al., 2019), it is possible that *Tgfb1* is directly or indirectly induced by CSF1. Indeed, it is highly-expressed by mouse bone marrow-derived and thioglycolate-elicited macrophages (Biogps.org). The unexpected macrophage αSMA expression in response to CSF1-Fc treatment may indicate TGFB signaling in macrophages. αSMA+ macrophages were described previously in a foreign body response (Mooney et al., 2010), and αSMA-expressing macrophages have also been implicated in protecting the bone marrow environment from radiation-induced injury (Ludin et al., 2012). Hence, in the context of tissue injury, αSMA+ cannot be considered a myofibroblast marker. It is also possible that *Tgfb1* is induced as a response to rapid liver growth, to limit expansion, given its known anti-proliferative function (Michalopoulos and Bhushan, 2021).

The function of HPC activation in liver fibrogenesis vs regeneration is controversial (Michalopoulos and Bhushan, 2021; Williams et al., 2014). Partial hepatectomy in fibrotic liver was previously reported to drive pro-fibrogenic HPC activation that impaired liver regeneration (Kuramitsu et al., 2013). On the other hand, adoptive transfer of CSF1-stimulated macrophages (BMDM) that reduced fibrosis in the CCL4 model was associated with TWEAK-dependent HPC activation (Thomas et al., 2011), and even transfer of BMDM into healthy mice induced transient macrophage TWEAK-dependent HPC activation (Bird et al., 2013). In the TAA model, HPC activation resolved spontaneously whereas fibrosis did not, and CSF1-Fc treatment had no effect in either healthy or TAA-exposed mice. Hence, we do not provide support for direct engagement of recruited macrophages with HPC in this setting.

Although our results are promising, some impacts of CSF1-Fc treatment could produce a dose-limiting toxicity in CLD patients and would need to be monitored. Splenomegaly and thrombocytopenia occur in advanced liver disease and are associated with poor outcomes. Splenectomy improves liver function in patients with advanced liver disease, and also reduces fibrosis and augments liver function in mouse models (Yada et al., 2015; Zheng et al., 2020). Mechanistically, the spleen could be a source of circulating TGFB1 and the monocyte chemokine CCL2 (Li et al., 2018). Thrombocytopenia in CLD is multi-factorial, including platelet sequestration, reduced production and increased destruction (Mitchell et al., 2016). Transient thrombocytopenia was the dose-limiting toxicity in initial human clinical trials of CSF1(Garnick and O’Reilly, 1989). This was further investigated in mice and found to be independent of the spleen and platelet production, and to resolve with prolonged treatment (Baker and Levin, 1998). The impact of CSF1 on platelets was rather attributed to increased activity of monocytes/macrophages which shortened platelet survival, but subsequently increased platelet production compensated for ongoing destruction (Baker and Levin, 1998). In the current study, we observed transient thrombocytopenia, even with low-dose CSF1-Fc, but this was rapidly resolved.

In conclusion, strategies to ‘reprogram’ macrophages have significant therapeutic potential via stimulation of multiple coordinated pro-regenerative macrophage functions, including phagocytosis, matrix remodeling, angiogenesis and production of tissue trophic factors. Here we have demonstrated striking impacts of CSF1-Fc on fibrosis and regrowth of fibrotic liver post-PHx. The therapeutic impacts may be attributable to CSF1 signaling specifically and/or driven by the increase in liver macrophages and amplified within the tissue microenvironment. The timing of intervention is critical because macrophages are shaped by the evolving microenvironment at the site of injury, as clearly illustrated by the different outcomes of CSF1-Fc treatment during and after cessation of liver injury. Further delineation of the molecular programs that drive restorative macrophage activities at the expense of their pathological functions may uncover other novel macrophage reprogramming strategies that could be harnessed to reduce the global burden of chronic liver disease.

## Materials and Methods

### CSF1-Fc reagents

This study utilised two CSF1-Fc reagents with equivalent biological impacts. The original porcine (P)-CSF1-Fc reagent was utilised at 1 mg/kg as in previous reports (Gow et al., 2014; Irvine et al., 2020; Sauter et al., 2016). The novel human CSF1-mouse Fc conjugate (HM-CSF1-Fc) was utilised at 5 mg/kg, as this dose elicited increased circulating monocytes, liver and spleen weight similar to 1 mg/kg P-CSF1-Fc. 1 mg/kg HM-CSF1-Fc was used as a sub-maximal dose in the setting of partial hepatectomy (**Fig. 7–8**), as this dose induced liver growth without monocytosis (**Fig. S2A-C**). Pig and human CSF1 proteins are both active on mouse CSF1R (Gow et al., 2012) and in our hands the Fc conjugates have similar activity on mouse bone marrow (not shown). The difference in efficacy may reflect different pharmacokinetics. Pig immunoglobulin does not bind to human Fc receptors (Shields et al., 2001) and the pig IgG1A Fc fragment used (Gow et al., 2014) is completely divergent in the critical FcR binding domain defined by site-directed mutagenesis (Egli et al., 2019) that is shared by mouse and human immunoglobulins. The mouse Fc domain used herein has the L234A/L235A mutations that reduce but do not abolish binding to mouse FcR and to C1q (Arduin et al., 2015).

### Animals

Studies were approved by a University of Queensland animal ethics committee. 6-8 week old C57Bl6/J mice were sourced from the Animal Resource Centre (Perth, Australia) and housed in a specific pathogen-free facility. Animals were randomly assigned to CSF1-Fc and saline treatment groups with mixed treatments in individual cages. For induction of liver fibrosis 300 mg/L thioacetamide (TAA, Sigma) was added to the sole source of drinking water. CSF1-Fc was administered by sub-cutaneous injection. At sacrifice blood was collected by cardiac puncture for hematology analysis (Mindray BC-5000) and serum separation (biochemical analysis by the University of Queensland Veterinary Laboratory Services).

### Partial hepatectomy (PHx)

Mice were anaesthetized by isoflurane inhalation. When fully anaesthetized the mouse was placed supine on a warming pad. A 1.5 cm upper midline incision was made. After the liver and the ligamentum falciforme were exposed the ligamentum was divided to the level of the superior vena cava to loosen the liver from the diaphragm. To achieve 50% PHx the left lateral and left median lobes were removed. The liver lobes to be resected were gently moved using saline-moistened cotton buds. A 5/0 suture was positioned around the appropriate lobe as near as possible to its base and tied with three knots. The lobe was removed distal to the suture leaving a short tissue stem. The resected lobes were retained and analysed as the pre-PHx baseline histology. Following surgery the abdominal cavity was rinsed with saline and the incision and skin were closed with a coated polyglactin 4/0 suture. The wound was disinfected, and the lost fluids were replaced by subcutaneous injection of up to 1 ml sterile saline. Mice were injected twice daily with buprenorphine for pain relief.

### Histology

Livers were fixed in 4% paraformaldehyde and paraffin-embedded. For immunostaining, epitope retrieval was performed in Diva Decloaker (Biocare Medical) followed by staining for Ki67 (Abcam ab16667, 1:100), F4/80 (Serotec MCA497GA, 1:400), wide-spectrum keratin (CKWSS, which labels bile duct epithelium and hepatic progenitor cells (Dako #Z0622, 1:400)) or SMA (Dako #M0851, 1:200). Secondary detection was with DAKO Envision HRP reagents or anti-species fluorophore conjugates (Thermofisher). Image quantification was performed from whole-slide digital images (VS120 scanner, Olympus) using ImageJ or Visiopharm software.

### Flow cytometry

Liver non-parenchymal cells were isolated as previously described (Melino et al., 2016). Briefly, tissue disaggregation was performed by finely chopping liver samples (~1-2 g) in 10 ml digestion solution containing 1 mg/ml Collagenase IV (Worthington) and 20 μg/ml DNAse1 (Roche) and incubating at 37°C for 45 min on a rocking platform prior to mashing through a 70 uM filter (Falcon). The cell pellet was collected by centrifugation and resuspended in an isotonic 30% Percoll solution to separate hepatocytes and non-parenchymal cells. Cells were stained for a panel of myeloid markers (F4/80-AF647 (1:150), Cd11b-BV510 (1:200), Ly6G-BV785 (1:200), MHCII-BV421 (1:200), Tim4-PE-Cy7 (1:300), Ly6C-PE (1:300) (Biolegend) in buffer containing 2.4G2 supernatant to block Fc binding, washed and resuspended in buffer containing viability dye 7AAD (Life Technologies) for acquisition using a Cytoflex (Becton Dickinson). Live single cells were identified for phenotypic analysis by excluding doublets (FSC-A > FSC-H), 7AAD+ dead cells and debris. Single colour controls were used for compensation and unstained and fluorescence-minus-one controls were used to confirm gating. Data were analysed using FlowJo 10 (Tree Star). Cell counts were calculated by multiplying the frequency of the cell type of interest by the total mononuclear cell yield/gram of disaggregated tissue.

### qPCR

Liver samples were collected in TRIzol for RNA extraction and cDNA synthesis (Bioline) using standard protocols. RT-PCR was performed using the SYBR Select Master Mix (Thermofisher) on an Applied Biosystems QuantStudio system. Primer pairs used in this study are as follows: *Hprt* F: GCAGTACAGCCCCAAAATGG, *Hprt* R: AACAAAGTCTGGCCTGTATCCAA; *Tbp* F: CTCAGTTACAGGTGGCAGCA, *Tbp* R: ACCAACAATCACCAACAGCA; *Adgre1* F: CTGTCTGCTCAACCGTCAGGTA, *Adgre1* R: AGAAGTCTGGGAATGGGAGCTAA; *Ccl2* F: CAAGATGATCCCAATGAGTAGGC, *Ccl2* R: CTCTTGAGCTTGGTGACAAAAACTA; *Ccr2* F: GAACTTGAATCATCTGCAAAAACAAAT, *Ccr2* R: GGCAGGATCCAAGCTCCAAT; *Acta2* F: GATCCTGACTGAGCGTGGCTAT, *Acta2* R: CGTGGCCATCTCATTTTCAAAG; *Colla1* F: AGGGATCCAACGAGATCGAG, *Colla1* R:CAAGTTCCGGTGTGACTCGT; *Col3a1* F: TGGGATCAAATGAAGGCGAAT, *Col3a1* R: GCTCCATTCCCCAGTGTGTTTAG; *Mmp9* F: AGGGGCGTGTCTGGAGATTC, *Mmp9* R: TCCAGGGCACACCAGAGAAC; *Mmp13* F: ACAAAGATTATCCCCGCCTCAT, *Mmp13* R: GGCCCATTGAAAAAGTAGATATAGCC; *Plau* F: GGCTTTGGAAAAGAGTCTGAAAGTG, *Plau* R: GCCATAGTAGTGGGGCTGCAT; *Tgfb1* F: GTGGCTGAACCAAGGAGACG, *Tgfb1* R: GGCTGATCCCGTTGATTTCC; *Hbegf* F: CTGAGGAGGACCTGAGCTATAGGA, *Hbegf* R: GTTTTCATGGCTGCTGGTGA; *Hgf* F: ATTGGATCAGGACCATGTGAGG, *Hgf* R: CACATCCACGACCAGGAACA; *I16* F: AAATCGTGGAAATGAGAAAAGAGTTG, *I16* R: GCATCCATCATTTCTTTGTATCTCTG; *Tnf* F: GGTCCCCAAAGGGATGAGAAG, *Tnf* R: TCGAATTTTGAGAAGATGATCTGAGTG; *Brg1* F: GAAAGTGGCTCTGAAGAGGAGG, *Brg1* R: TCCACCTCAGAGACATCATCGC;

### Doppler Imaging

Hepatic blood flow was assessed by Power Doppler imaging using a Vevo 2100 ultrasound system fitted with a MS250 transducer (20 MHz center frequency (Fujifilm Visualsonics)). Scan settings were pulse repetition frequency (PRF) at 3 KHz, Doppler gain at 37 dB, medium persistence (frame averaging), and scan distance of ^~^20 mm, with a step size of 0.150 mm. Calculation of liver percent vascularity (PV), and 3D image reconstruction, were achieved using Vevolab analysis software v5.5.1.

### Data analysis

Sample sizes were determined by previous experiments utilizing CSF1-Fc treatment that detected statistically significant impacts on liver growth and regeneration in healthy mice and acute injury models, as well as our prior experience with the TAA model of liver fibrosis (Gow et al., 2014; Irvine et al., 2020; Irvine et al., 2015; Melino et al., 2016; Stutchfield et al., 2015). Analysis of histological and flow cytometry outcome data was performed blinded to treatment group. Data are presented as mean +/− standard deviation. Statistical tests were performed using GraphPad Prism 7.03. Data normality was tested using the Shapiro-Wilk test; unless otherwise stated, ordinary one-way ANOVA with Sidak’s multiple comparisons testing was used. All tests were two-tailed. All authors had access to the study data and reviewed and approved the final manuscript.

## Acknowledgements

KMI and DAH are grateful for core laboratory support from the Mater Foundation. We appreciate the support of the Preclinical Imaging, Biological Resources, Histology, Microscopy and Flow Cytometry Core Facilities at the Translational Research Institute.

